# Supervised contrastive learning enhances MHC-II peptide binding affinity prediction

**DOI:** 10.1101/2023.12.21.572942

**Authors:** Long-Chen Shen, Yan Liu, Zi Liu, Yumeng Zhang, Zhikang Wang, Yuming Guo, Jamie Rossjohn, Jiangning Song, Dong-Jun Yu

**Author notes:** E-mail Address: Long-Chen Shen.

## Abstract

Accurate prediction of major histocompatibility complex (MHC)-peptide binding affinity can improve our understanding of cellular immune responses and guide personalized immunotherapies. Nevertheless, the existing deep learning-based approaches for predicting MHC-II peptide interactions fall short of satisfactory performance and offer restricted model interpretability. In this study, we propose a novel deep neural network, termed ConBoTNet, to address the above issues by introducing the designed supervised contrastive learning and bottleneck transformer extractors. Specifically, the supervised contrastive learning pre-training enhances the model’s representative and generalizable capabilities on MHC-II peptides by pulling positive pairs closer and pushing negative pairs further in the feature space, while the bottleneck transformer module focuses on MHC-II peptide interactions to precisely identify binding cores and anchor positions in an unsupervised manner. Extensive experiments on benchmark datasets under 5-fold cross-validation, leave-one-molecule-out validation, independent testing, and binding core prediction settings highlighted the superiority of our proposed ConBoTNet over current state-of-the-art methods. Data distribution analysis in the latent feature space demonstrated that supervised contrastive learning can aggregate MHC-II-peptide samples with similar affinity labels and learn common features of similar affinity. Additionally, we interpreted the trained neural network by associating the attention weights with peptides and innovatively find both well-established and potential peptide motifs. This work not only introduces an innovative tool for accurately predicting MHC-II peptide affinity, but also provides new insights into a new paradigm for modeling essential biological interactions, advancing data-driven discovery in biomedicine.

## 1. Introduction

Major histocompatibility complex (MHC) molecules are a class of cell surface glycoproteins that play an essential role in the immune system(Holling, Schooten, & van Den Elsen, 2004; Tsai & Santamaria, 2013). They can be categorized into two major types: MHC class I (MHC-I) and MHC class II (MHC-II). Among them, MHC-II molecules can recognize exogenous antigens derived from pathogenic microorganisms or self-tissues to form MHC-peptide complexes, and then present them on the surface of antigen-presenting cells (APC) to CD4^+^ T cell receptors (TCR), thereby activating specific T cell immune responses and eliminating pathogens or damaged tissues (Sagan, et al., 2023). Accurate identification of MHC binding peptides is thus crucial not only for identifying neoepitopes that effectively trigger immune response targeting cancer or infected cells but also for providing key insights into vaccine design (Blass & Ott, 2021), immunotherapy (Wang, et al., 2020), and research on immune-related diseases (Liu, Shao, & Fu, 2021). In this context, computational methods can assist in identifying potential peptides for effective immune response, enhancing vaccine safety and efficacy, and guiding treatments for immune-related diseases such as autoimmune disorders, allergies, and cancer (Alspach, et al., 2019; Meraviglia-Crivelli, et al., 2022; Moore & Nishimura, 2020).

The MHC-II molecule is composed of the α and β chain. Each chain possesses a domain, specifically alpha-1 and beta-1, which collectively create a peptide-binding groove for the attachment of an antigenic peptide (Kisielow, Obermair, & Kopf, 2019). MHC-II molecules in humans are encoded by various Human Leukocyte Antigen (HLA) genes like HLA-DR, HLA-DQ and HLA-DP.(Traherne, 2008) Each MHC-II gene possesses numerous alleles, resulting in substantial genetic diversity within the population (Tadros, Eggenschwiler, Racle, & Gfeller, 2023). Unlike MHC-I, the open groove of MHC-II allows for hypervariable peptide lengths, usually between 10-30 amino acids, with a typical range of 13-17 residues (Chang, Ghosh, Kirschner, & Linderman, 2006). In essence, the polypeptides’ ends extend past the binding groove, with only some key amino acid residues (i.e., anchor residues) tightly coupled with MHC-II molecules, ensuring the stability of protein-peptide interaction (Jones, Fugger, Strominger, & Siebold, 2006). This characteristic allows MHC-II molecules to effectively present exogenous antigens to CD4^+^ T cells, thus regulating immune responses(Couture, et al., 2019; Neefjes, Jongsma, Paul, & Bakke, 2011). However, experimental characterization of the binding specificity with numerous MHC-II molecules is time-consuming and laborious. In this regard, computational methods can be employed as effective alternatives to quantify peptide-MHC-II molecule binding affinity. Recent studies indicate the growing importance of such computational methods and tools in the fields of vaccine design and immunotherapy (Fatima, et al., 2022; Finotello, Rieder, Hackl, & Trajanoski, 2019; Soleymani, Tavassoli, & Housaindokht, 2022).

MHC-II peptide binding prediction methodologies mainly comprise allele-specific and pan-specific methods. While allele-specific methods are restricted to predicting binding affinity for MHC-II molecules found in the training set, pan-specific methods can predict unseen MHC-II molecules with known protein sequences (Jensen, et al., 2018). Pan-specific methods have broader applications and encompass allele-specific methods. The classic allele-specific method, NetMHCII (Jensen, et al., 2018), trained separate artificial neural networks (ANNs) for each MHC-II molecule to predict peptide-binding affinity. Given the scarcity of experimental data for most MHC-II molecules and the limited scope of allele-specific methods, pan-specific approaches emerged for MHC-II peptide binding prediction (You, Qu, Mamitsuka, & Zhu, 2022; Zeng & Gifford, 2019). NetMHCIIpan-3.2 (Jensen, et al., 2018) was the first pan-specific method to predict binding affinity using ANN. It combines MHC-II pseudo-sequence information and integrates 40 networks with different numbers of hidden neurons. Zeng *et al*. proposed a multi-model ensemble method PUFFIN (Zeng & Gifford, 2019) that quantified prediction uncertainty and prioritized peptides with “binding likelihood” to improve the accuracy of high-affinity peptide selection. MHCAttnNet (Venkatesh, Grover, Srinivasaraghavan, & Rao, 2020) incorporated variable-length peptide sequences and the MHC allele via attention mechanism and Bi-LSTM encoder, thus improving MHC-I and II peptide binding predictions. Similarly, DeepSeqPanII (Z. Liu, et al., 2021), a recurrent neural network model with attention mechanism, also predicted peptide-HLA class II binding. Recently, You *et al*. (You, et al., 2022) proposed a binding core-aware interaction model to enhance MHC-II peptide binding affinity prediction. Despite impressive advances in prediction performance, current methods have some drawbacks: (1) the large diversity of MHC-II molecules and scarcity of binding data often lead to low prediction performance with limited training data; (2) while some algorithms aim to enhance model interpretability by analyzing attention modules, there is still much room for improvement; (3) most methods singularly characterize MHC-II molecules and peptide sequences, which may lead to insufficient mining of sequence information; (4) Most algorithms currently ensemble multiple models of similar structure to boost model performance. However, it will save computing resources for training and prediction if comparable or superior performance can be achieved with fewer integrations or a single model.

Here, we present ConBoTNet (<u>Con</u>trastive <u>Bo</u>ttleneck <u>T</u>ransformer <u>Net</u>work), a new deep-learning framework that employs powerful supervised contrastive learning with the bottleneck transformer extractor to enable accurate prediction of MHC-II peptide binding affinity. In the supervised contrastive learning pre-training phase, the continuous values of binding affinity are divided into ordered categories, and then the supervised contrastive loss function is applied to minimize the normalized embeddings within each class and maximizes the distances between different classes. Moreover, the bottleneck transformer modules can detect the binding core and precise anchor positions from MHC-II molecules and peptide interaction features due to their robustness in feature extraction. Performance evaluations, including 5-fold cross-validation (5-fold CV), leave-one-molecule-out (LOMO) validation, and independent testing on benchmark datasets, reveal that our model outperforms five current state-of-the-art (SOTA) methods, exhibiting rapid training convergence and impressive generalization capacity.

## 2. Materials and methods

### 2.1. Benchmark datasets

In this study, three public benchmark datasets (BD2016, BC2015, BD2023) were used as benchmark datasets to evaluate and compare the performance with other competing methods. The sources and construction process of these three datasets are introduced in this section.

#### BD2016

This dataset includes the MHC-II peptide binding affinity data extracted from the IEDB database (Vita, et al., 2019) (https://www.iedb.org/) in 2016 by NetMHCIIpan-3.2 (Jensen, et al., 2018). The binding values (*IC*_50_) of the experimental data were normalized to the range of [0,1] by 1 − log (*IC*_50_ *nM*)/log (50,000). The BD2016 dataset encompasses 134,281 MHC-II peptide affinity data for 36 HLA-DR, 27 HLA-DQ, 9 HLA-DP alleles, and 8 H-2 alleles, respectively. It is considered as the gold standard dataset by a number of previous affinity prediction methods, providing widely accepted 5-fold CV data partitioning by grouping the peptides with common motifs into the same fold (Z. Liu, et al., 2021; You, et al., 2022; Zeng & Gifford, 2019). This dataset is accessible at https://services.healthtech.dtu.dk/services/NetMHCIIpan-3.2/.

#### BC2015

This dataset included 51 MHC-II peptide complexes with crystal structures and has been used as the benchmark dataset by several MHC-II peptide affinity prediction algorithms, particularly for unsupervised binding core evaluations. It is also accessible at https://services.healthtech.dtu.dk/services/NetMHCIIpan-3.2/.

#### BD2023

an up-to-date independent test set curated from an automatically updated MHC-II binding prediction platform (http://tools.iedb.org/auto_bench/mhcii/weekly/) (Andreatta, et al., 2018). Specifically, we selected recently added MHC-II peptide binding data generated from 2022-04-01 to 2023-04-21 and removed the duplicates from the training set BD2016. We then applied the filtering criterion to only include MHC-II molecules with demonstrated binding to a minimum of 28 peptides, to ensure that the dataset could focus on significant binding interactions. The dataset contained samples of two measurement types, i.e., *IC*_50_ and binary (binding or decoy) labels. Performance evaluation metrics used by the experiments are provided in the **Supplementary Text S1**.

### 2.2. Overview

The deep-learning framework of ConBoTNet is illustrated in **Figure 1**, which mainly includes the following modules: (1) sequence encoding for peptides and MHC-II molecules; (2) backbone encoder network for feature extraction; (3) projection network used for contrastive learning; (4) binding core and affinity predictor. The core contributions of ConBoTNet are as follows: (i) Combining supervised contrastive learning with a deep transformer-based network, we could effectively learn the latent representations of the MHC-II peptide binding complex. (ii) Then, we can smoothly adapt the pre-trained model to various downstream tasks, including binding affinity prediction and binding core identification.

**Figure 1.**
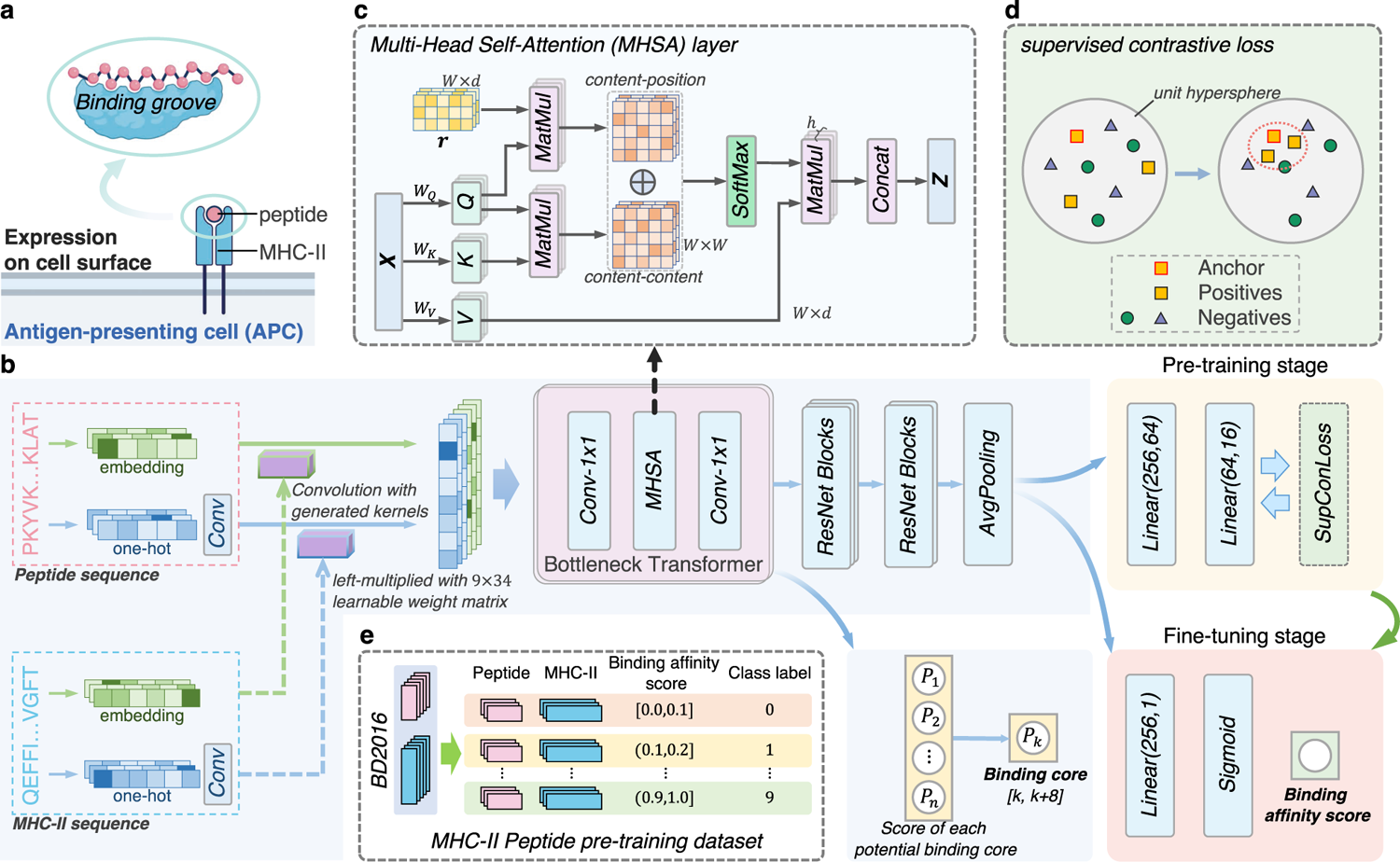
Model architecture of ConBoTNet. a, Schematic diagram of the binding of MHC-II molecules and peptides on the surface of the antigen-presenting cell. b, Architecture of ConBoTNet and the training pipeline. c, Detailed structure of the multi-head self-attention (MHSA) layer. d, Schematic diagram of the supervised contrastive loss. e, Based on the division of binding affinity values of BD2016, the data processing workflow for constructing the pre-training classification dataset.

### 2.3. Sequence encoding

The input to ConBoTNet includes the peptide and MHC-II molecule sequence pairs, whose binding schematic is illustrated in **Figure 1a**. In particular, the MHC-II molecule sequence is represented by a pseudo-sequence comprising 34 amino acids in contact with the peptide-binding core (distance within 4Å) (Karosiene, et al., 2013). The first 15 amino acids of these are from the α-chain while the rest are from the β-chain, following the strategy used in most previous studies (Holland, Cole, & Godkin, 2013; Jensen, et al., 2018; You, et al., 2022; Zeng & Gifford, 2019).

We define *P* as the input peptide sequence with a length of *L* and *Q* as the MHC-II pseudo-sequence with a length of 34, respectively. To capture the hidden semantic information and ensure the complete representation of the input information, we encode both the peptide sequence and the MHC-II pseudo-sequence using the embedding layer and one-hot encoding.

For the embedding layer, let *d* be the dimension of amino acid embeddings, *X*_e_ ∈ ℝ^L×d^, the output of the embedding layer for peptide sequence *P*, and *Y*_e_ ∈ ℝ^34×d^, the output of the embedding layer for the MHC-II pseudo-sequence *Q*, are given as follows:

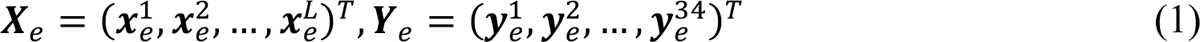

where *x*_e_*^i^* ∈ ℝ^d^ and *y*_e_^j^ ∈ ℝ^d^ are the feature vectors of the *i*th amino acid of *P* and *j*th amino acid of *Q*, respectively.

In terms of one-hot encoding, *d*^’^=20 is the dimension of one-hot encoding, representing 20 common amino acids. *X* ∈ ℝ^L×d^, the one-hot encoding for peptide sequence *P*, and *Y* ∈ ℝ^34×d^, the one-hot encoding for MHC-II pseudo-sequence *Q*, are given as follows:

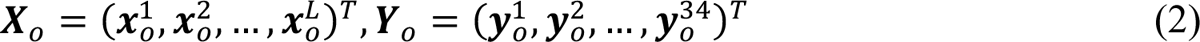

where *x*_o_^i^ ∈ ℝ^d’^ and *y*_o_^j^ ∈ ℝ^d’^ are the one-hot feature vectors of the *i*th amino acid of *P* and *j*th amino acid of *Q*, respectively. A value of 1 was assigned to the corresponding amino acid position in the input sequences and 0 elsewhere.

### 2.4. Backbone encoder network

Next, we briefly introduce the architecture of ConBoTNet in combination with **Figure 1b**. Firstly, we used two feature encoding methods to characterize the peptide sequence and the MHC-II molecular pseudo-sequence, respectively. Subsequently, we built the interaction features through the interaction module proposed by DeepMHCII (You, et al., 2022). It is noteworthy that we added 64 1 × 3 convolution filters after one-hot encoded features for peptide and MHC-II, respectively. We then combined the interaction features from different encoding methods as the final peptide MHC-II molecule interaction features. Next, these were sequentially passed through two bottleneck transformers, several one-dimensional ResNet blocks, and an avg-pooling layer.

We refer to the above pipeline as the backbone encoder network, denoted as *Enc*(⋅), which is expressed by the following formula:

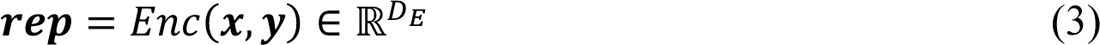

We define the number of samples in a batch as *N*. *x*, *y* represent pairs of peptide and MHC-II molecules. The encoder network map *x*, *y* to a representation vector *rep*, which is normalized to the unit hypersphere in ℝ^DE^ (*D*_E_ = 256, according to the complexity of features).

#### The details of Bottleneck transformers

The current state-of-the-art methods for predicting MHC-II peptide binding utilize shallow or deep convolutional networks, such as PUFFIN (Zeng & Gifford, 2019), DeepMHCII (You, et al., 2022). However, we propose that incorporating the transformer module can further enhance the predictions. Transformers capture long-term dependencies in MHC-II peptide binding features and enhance context awareness through multi-head self-attention, enabling them to learn the universal pattern and contextual information of MHC-II peptide binding. Inspired by the bottleneck transformer (BoTNet)(Srinivas, et al., 2021) architecture and its outstanding performance in computer vision tasks, we integrated it with the domain knowledge of affinity prediction, and improved its Multi-head Self-Attention (MHSA) layer to capture the contextual and global information related to MHC-II peptide interaction.

We illustrated an MHSA layer in **Figure 1c**. The multi-head self-attention layer can be summarized by the following equation:

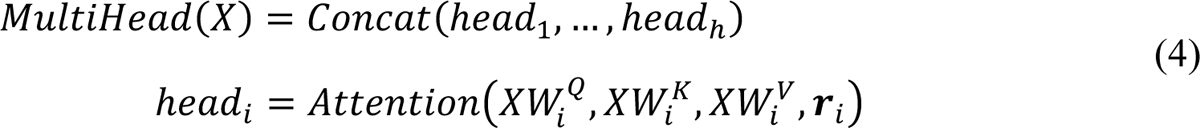

where *W*_i_^Q^ ∈ ℝ^dmodel x^ dk, *W*_i_^K^ ∈ ℝ^dmodel x dk^, *W*_i_^v^ ∈ ℝ^dmodel x dk^ represents the parameter matrices, *r*_i_ ∈ ℝ^dk×dw^ is the relative position encodings of interaction features, *d*_w_ is the length of features and *i* denotes the *i*-th attention head. *d*_*model*_ is the input feature dimension of the MHSA layer. In this work, we use *d*_model_ = 128, ℎ = 4, and *d*_K_ = *d*_model_/ℎ = 32.

As shown in the MHSA layer in **Figure 1c**, the input size is *W* × *d*, which respectively represent the length and encoding dimension of the MHC-II peptide interaction feature map. *r*_*i*_ denote a learnable relative position encodings matrix of the *head*_i_, *r* is the set of all *r*_i_. First, the input feature *X* is multiplied by the weight matrices *W*_3_, *W*_4_ and *W*_5_ respectively to obtain matrices *q*, *k* and *v*. Then, matrix *r* is multiplied by *q* to get the *content-position* which forms part of the attention. Following traditional self-attention mechanism, the product of *q* and *K* is calculated as *content-content*. By adding corresponding elements from the *content-position* and *content-content*, and then applying the softmax function, we calculate the final attention weights. Finally, these attention weights are multiplied by the matrix *v* to obtain the output feature map of the self-attention layer with the position information.

### 2.5. Supervised contrastive learning

#### Data preprocessing for supervised contrastive pre-training

In the pre-training phase of supervised contrastive learning, we split the training dataset, BD2016, into ten distinct categories at intervals of 0.1 based on the transformed affinity score. **Figure 1e** provides a detailed illustration of this categorization — for instance, category 0 includes affinity values within the range of [0.0, 0.1]. We examined the category numbers such as 5, 8, 10, and 14. As a result, 5-fold CV test confirmed 10 as the best choice, as can be seen from **Table S1**. Particularly, having a high number of categories led to fewer samples per group, which hindered the learning of discriminative features. Conversely, when there were too few categories available, it would prevent the full realization of the substantial potential in supervised contrastive learning. Therefore, we finally chose to retain the categorization number of ten.

#### Supervised contrastive learning pre-training

Next, we briefly introduce the main components of the supervised contrastive learning framework and the expression equation of the supervised contrastive loss function.

We mainly divide the supervised contrastive learning framework into two parts: Encoder Network and Projection Network. The specific architecture of the encoder network is described in detail in ***backbone encoder network*** section. We represent the output features of the encoder network as *rep*. Following the encoding network, we employ a multi-layer projection network to map *rep* to vector *z*. The projection network is represented by the following formula:

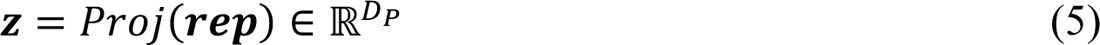

where *Proj*(⋅) consists of two fully-connected layers and a ReLU activation function subsequently normalizing the final output to the unit hypersphere in ℝ^0*^ (where *D*_E_ = 16 in this study). In the end, we utilize the inner product between *z* as a metric for their distance in the projected space, and calibrate the model parameters via the contrastive loss, which is expressed as follows:

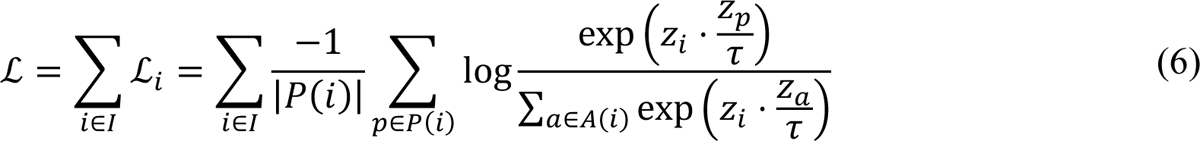

In a given batch, let {(*x*_K_, *y*_K_), *label*_K_}_KC),*,…,F_ be peptide, MHC-II, label triplets and *z*_9_ = *Proj*(*Enc*(*x*_K_, *y*_K_)) ∈ ℝ^0*^. Let *i* ∈ *I* ≡ {1,2, …, *N*} be the index of a sample, typically referred to as the ‘anchor’. *A*(*i*) ≡ *I*\{*i*} represents the set of indices not included in set *I*, *P*(*i*) ≡ {P ∈ *A*(*i*): *label*_b_ = *label*_i_} is the set of indices for all positives in the batch excluding *i*, and |*P*(*i*)| is its cardinality. The ⋅ stands for the inner (dot) product, while τ ∈ ℝ^G^ is a scalar representing a temperature parameter. A diagram of the supervised contrastive loss is shown in **Figure 1d**, where the distance of MHC-peptide pairs within the anchor category is pulled closer.

Upon completing the supervised contrastive learning phase, we discard the projection network *Proj*(⋅). In the subsequent fine-tuning stage, we employ the Mean Squared Error (MSE) loss function to perform further model training based on the established encoder *Enc*(⋅) parameters.

### 2.6. Two stages training process

In the pre-training stage, we mapped the interaction feature *rep*, with a dimension of 256, to a representation vector *z*, with a dimension of 16, via the projection network *Proj*(⋅). We optimized the model parameters with the supervised contrastive loss and larger batch sizes (set to 512) for model stability and speed. In the fine-tuning stage, we replaced the projection network with a linear layer and sigmoid function, then optimized the pre-trained model with MSE loss. Additionally, we determined the binding cores by identifying the highest-scoring positions from the output vector of the bottleneck transformer during inference. A detailed description of the model implementation and hyperparameter settings is provided in **Supplementary Text S2**.

## 3. Results

### 3.1. Comparison of ConBoTNet and competing methods under 5-fold cross-validation

**Table 1** presents the average area under the ROC curve (AUC) and Pearson correlation coefficient (PCC) of ConBoTNet and competing methods over all MHC-II molecules via 5-fold CV on BD2016 data set. Since NetMHCIIpan-4.0 has already incorporated elution ligand mass spectrometry(Purcell, Ramarathinam, & Ternette, 2019) data, it is not suitable to compare NetMHCIIpan-4.0 through 5-fold CV and LOMO. Our proposed model, ConBoTNet, outperforms existing models in terms of both average AUC and PCC. Specifically, ConBoTNet outperformed the existing SOTA method, DeepMHCII, with an AUC of 0.865 (95% confidence interval [CI]: 0.857-0.871) and a PCC of 0.709 (95% CI: 0.698-0.719), compared to the AUC of 0.856 achieved by DeepMHCII (95% CI: 0.847-0.864) and PCC of 0.690 (95% CI: 0.679-0.700), respectively. Moreover, the boxplots in **Figure 2a** clearly illustrate the overall performance of each method across 61 MHC-II molecules, where ConBoTNet outperforms all other methods. **Figure 2b** provides a comparison between ConBoTNet and the current state-of-the-art method, DeepMHCII, in terms of AUC and PCC values for each MHC-II molecule. Each dot in **Figure 2b** corresponds to one MHC-II molecule. Specifically, the scatter plots reveal that ConBoTNet exhibited varying degrees of performance improvement over 57 out of 61 MHC-II molecules (representing ∼93.4% of molecules) compared to DeepMHCII in terms of AUC and PCC.

**Figure 2.**
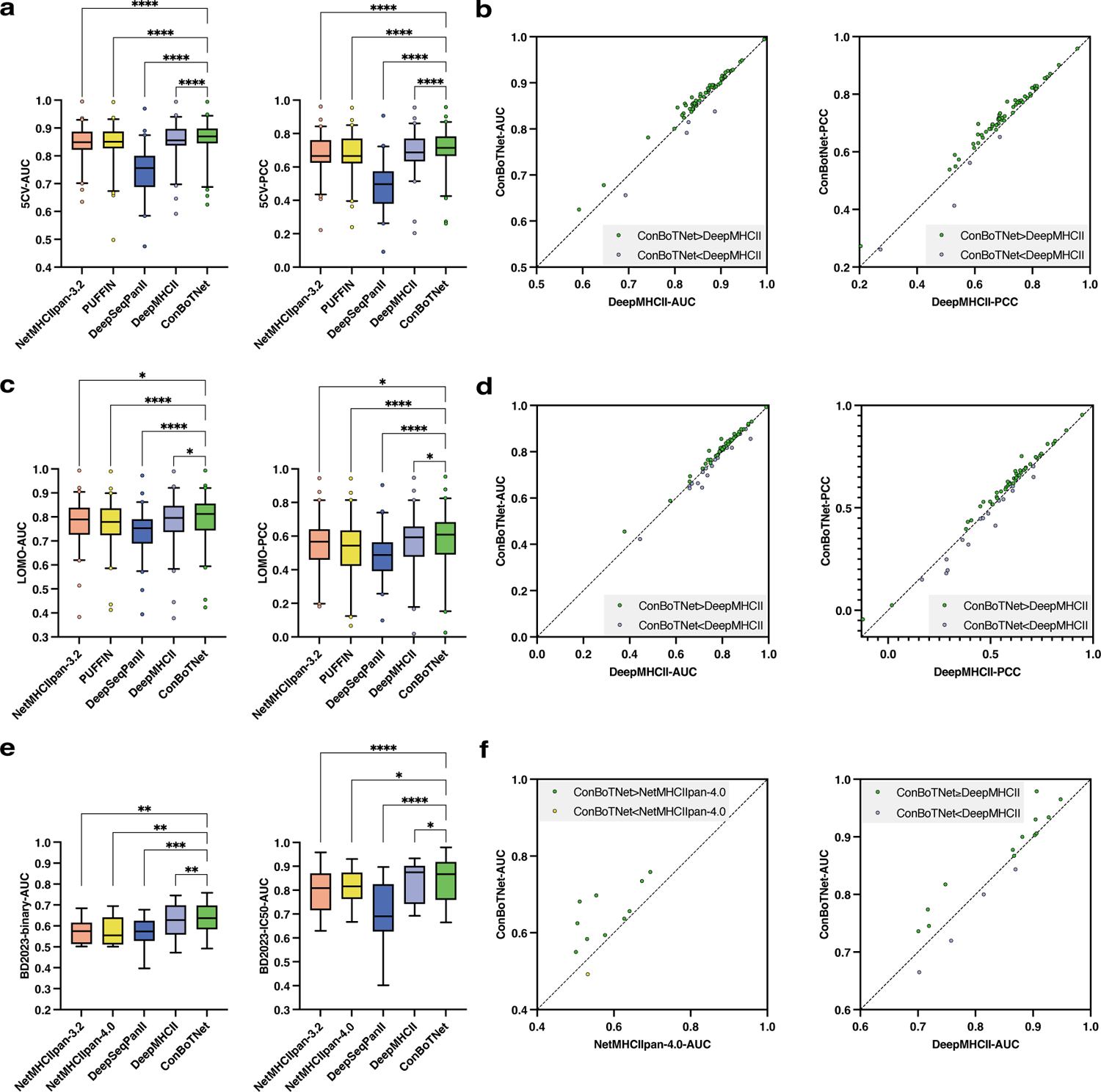
Performance evaluation of ConBoTNet and competing methods on different benchmarks. a, Boxplots of the performance of ConBoTNet and competing methods under 5-fold CV. b, Two scatter plots of point-to-point comparisons between ConBoTNet and the current best competing method on 5-fold CV under the AUC and PCC metrics, respectively. c, Boxplots of ConBoTNet and competing methods on LOMO. d, Two scatter plots of point-to-point comparisons between ConBoTNet and the current best competing method on LOMO under the AUC and PCC metrics, respectively. e, Boxplots of the performance of ConBoTNet method and competing methods on BD2023 data set. f, Two scatter plots of point-to-point comparisons between ConBoTNet and the current best competing method on the BD2023 subset binary and *IC*_50_under the AUC index, respectively. Asterisks indicate statistical significance levels against other methods (*, *P*<0.05; **, *P*<0.01; ***, *P*<0.001; ****, *P*<0.0001).

**Table 1.** Performance comparison of ConBoTNet and competing methods.

| Method | 5-CV | | LOMO | | BD2023 (*binary*) | BD2023 ($IC_{50}$) | | Binding core |
| --- | --- | --- | --- | --- | --- | --- | --- | --- |
|  | AUC | PCC | AUC | PCC | AUC | AUC | PCC | No. of correct/No. of total |
| NetMHCIIpan-3.2 | 0.847 | 0.679 | 0.775 | 0.544 | 0.571 | 0.802 | 0.616 | **45/51** |
| NetMHCIIpan-4.0 | -<sup>a</sup> | - | - | - | 0.577 | 0.814 | 0.653 | - |
| PUFFIN | 0.846 | 0.676 | 0.768 | 0.525 | - | - | - | - |
| DeepSeqPanII | 0.741 | 0.488 | 0.739 | 0.485 | 0.569 | 0.704 | 0.416 | 10/51 |
| DeepMHCII | 0.856 | 0.690 | 0.783 | 0.558 | 0.620 | 0.831 | 0.674 | 44/51 |
| ConBoTNet | **0.865** | **0.709** | **0.789** | **0.568** | **0.637** | **0.845** | **0.682** | **45/51** |

<sup>a</sup> '-' indicates that the corresponding value does not exist.

### 3.2. Comparison of ConBoTNet and competing methods under LOMO

**Table 1** also shows that ConBoTNet transcended other competing methods in terms of average AUC and PCC under LOMO on BD2016 data set. The experimental results are consistent with 5-fold CV. Specifically, ConBoTNet achieved the highest AUC score of 0.789 (95% CI: 0.778-0.796) and PCC score of 0.568 (95% CI: 0.557-0.578), respectively. Furthermore, the boxplots in **Figure 2c** demonstrate that ConBoTNet outperforms competing methods in most cases. ConBoTNet shows statistically significant improvement over the suboptimal method, DeepMHCII (P-value = 0.0413). Additionally, **Figure 2d** illustrates that ConBoTNet outperforms the suboptimal method for approximately 72.1% and 73.8% of the MHC-II molecules based on the AUC and PCC metrics, respectively.

### 3.3. Comparison of ConBoTNet and competing methods on independent testing set

The BD2023 dataset was collected from the latest IEDB release by the MHC-II automated benchmark platform. Any overlapping items with BD2016 within BD2023 were subsequently removed. We then divided this dataset into two subsets: BD2023 (binary) and BD2023 (*IC*_50_), based on the types of measurement data. **Table 1** shows the evaluation results of each model on BD2023. For NetMHCIIpan-3.2 and NetMHCIIpan-4.0, these results were obtained from the benchmark testing platform. Meanwhile, the results for DeepSeqPanII, DeepMHCII, and ConBoTNet were obtained by training these models on BD2016 and then tested on this independent dataset. Unfortunately, we could not provide test results on BD2023 for the PUFFIN method, as the training code was not disclosed, and the author-provided model download link was invalid. According to **Table 1**, our model has shown improvements in terms of average AUC compared to the current best-performance models. Specifically, there was a 2.7% improvement in BD2023 (binary), with the AUC increasing from 0.620 (95% CI: 0.602-0.639) to 0.637 (95% CI: 0.618-0.657). For BD2023 (*IC*_50_), 1.7% improvement was achieved, with the AUC increased from 0.831 (95% CI: 0.795-0.865) to 0.845 (95% CI: 0.813-0.880). The distributions of AUC scores for each MHC-II molecules on both subsets are further depicted in **Figure 2e**, where ConBoTNet demonstrates statistically significant improvements compared to other models (paired t-test, P-value = 0.0095 vs. NetMHCIIpan-4.0 and P-value = 0.0467 vs. DeepMHCII). Additionally, **Figure 2f** shows that ConBoTNet outperforms the suboptimal methods (NetMHCIIpan-4.0 and DeepMHCII) for approximately 91.0% and 76.5% of the MHC-II molecules in the two BD2023 subsets, respectively. Detailed performance comparison results on the updated BD2024 independent test set are provided in the **Supplementary Text S3** and shown in **Figure S1**.

### 3.4. Comparison of ConBoTNet and single model based on ensemble methods

Various algorithms can enhance binding affinity prediction by ensembling multiple models of the same or similar architecture. For example, NetMHCIIpan-3.2 uses 40 neural networks for ensemble; PUFFIN is a deep ensemble model composed of 20 deep residual convolutional neural networks; DeepMHCII is an ensemble of 20 models using a simple average method. In this section, to examine the possibility of performance improvement by relying solely on the model architecture itself, we discuss the comparison between ConBoTNet and a single model of existing ensemble algorithms. However, we did not include PUFFIN in the performance comparison since it did not provide neither implementation code nor the trained model. We compared ConBoTNet with the SOTA affinity prediction method, DeepMHCII, which outperformed PUFFIN (discussed in detail in DeepMHCII by You *et al*.), and DeepSeqPanII, a single-model algorithm. The performance comparison results on 5-fold CV test for 61 MHC-II molecules are shown in **Figure 3**, while the overall performance evaluation metrics are presented in **Table S2**. As can be seen, ConBoTNet outperformed DeepMHCII on about 95.1% of MHC-II molecules with statistical significance (P-value < 0.0001). Specifically, ConBoTNet achieved the best average AUC of 0.837 on 5-fold CV, which was 1.9% higher than the SOTA method, DeepMHCII (0.821), and 13.0% higher than DeepSeqPanII (0.741), respectively. Similarly, the average PCC was also improved compared to DeepMHCII (increased by 4.6%, 0.657 versus 0.628), indicating the excellent performance of the ConBoTNet model. Moreover, a more detailed description of the ablation studies involving the impact of bottleneck transformer modules, supervised contrastive learning, as well as sequence similarity on the model performance is provided in the **Supplementary Text S4**.

**Figure 3.**
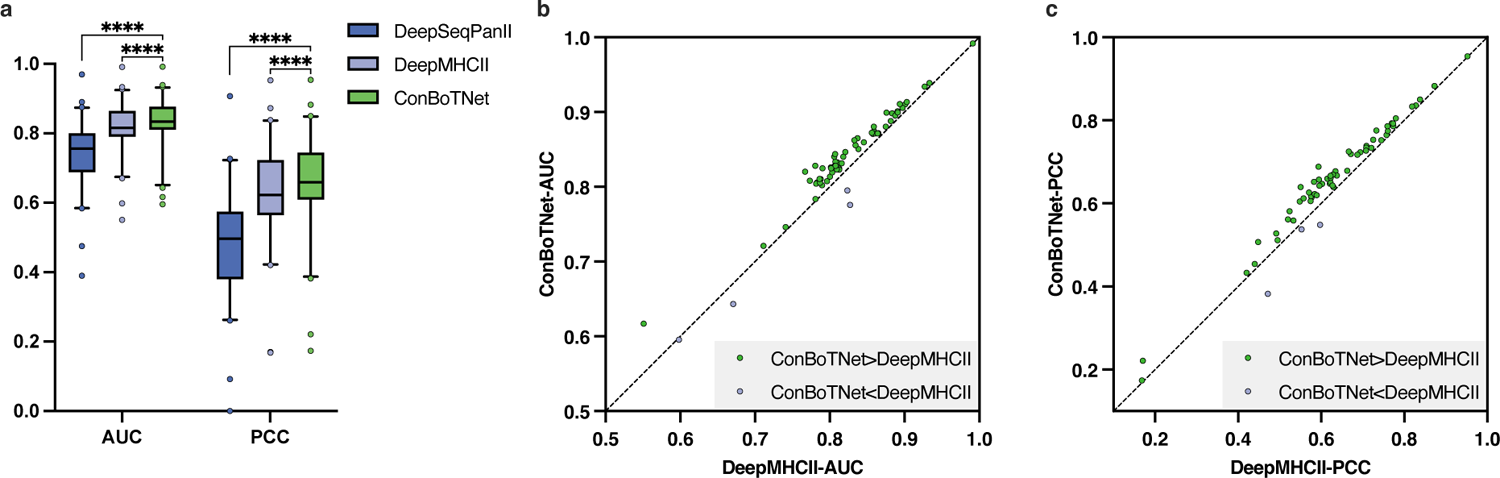
Performance evaluation of ConBoTNet and competing methods on a single model under 5-fold CV. a. Boxplot representing the performance of ConBoTNet, DeepSeqPanII, and DeepMHCII on 5-fold CV under the AUC and PCC metrics. b, c, The scatter plots of point-to-point comparisons between ConBoTNet and the current best competing method on 5-fold CV under the AUC and PCC metrics, respectively.

### 3.5. Binding core prediction & antigen weight visualization

The final column in **Table 1** shows the ability of ConBoTNet and the compared methods to predict the binding core on BC2015. It suggests that both ConBoTNet and NetMHCIIpan-3.2 achieved remarkable prediction accuracy, correctly identifying 45 out of 51 MHC-II peptide complexes. **Figure 4a** provides a detailed visualization of the results generated by ConBoTNet. Accurate predictions are denoted in blue, while incorrect predictions are highlighted in red. To better understand the model’s learning capability and potential error sources, we utilized the Captum (Kokhlikyan, et al., 2020) tool to analyze the attention weights assigned by the ConBoTNet model to different positions of the peptide sequences. These results are visualized through a heatmap in **Figure 4a**. As the binding core consists of a continuous fragment of nine amino acids within the peptide sequence, we demarcate this fragment using a box encompassing nine amino acids with the highest sum of interest. The accurately and inaccurately predicted binding cores are marked by blue boxes and red boxes, respectively. Excitingly, ConBoTNet demonstrates a strong focus on the most critical aspect of peptide binding affinity, i.e., the binding core. For some MHC-peptide complexes, such as PDB IDs 1UVQ and 1IAO, the ConBoTNet model may have failed to identify the real binding cores due to the influence of the peptides’ flanking regions (Holland, et al., 2013). Nevertheless, this observation underscores the model’s capability to emphasize the peptide fragment that is most important for the affinity effect. In **Figure 4b**, we present five PDB structures depicting the peptide-binding grooves of human MHC-II (HLA-DR, DP, DQ) and mouse MHC-II (H-2) molecules. Amino acids with the highest attention weights are usually located at the MHC peptide-binding pockets and involved in the polar interactions between the peptide and MHC-II molecule or TCR, e.g. In the HLA-DR2b/MBP-peptide complex (PDB ID: 1YMM), a bulky amino acid F8 of the peptide forms hydrogen bonds with Q9 and N62 at the MHC-II α chain. This also illustrates that our proposed ConBoTNet can identify binding anchors that interact with MHC-II or TCR.

**Figure 4.**
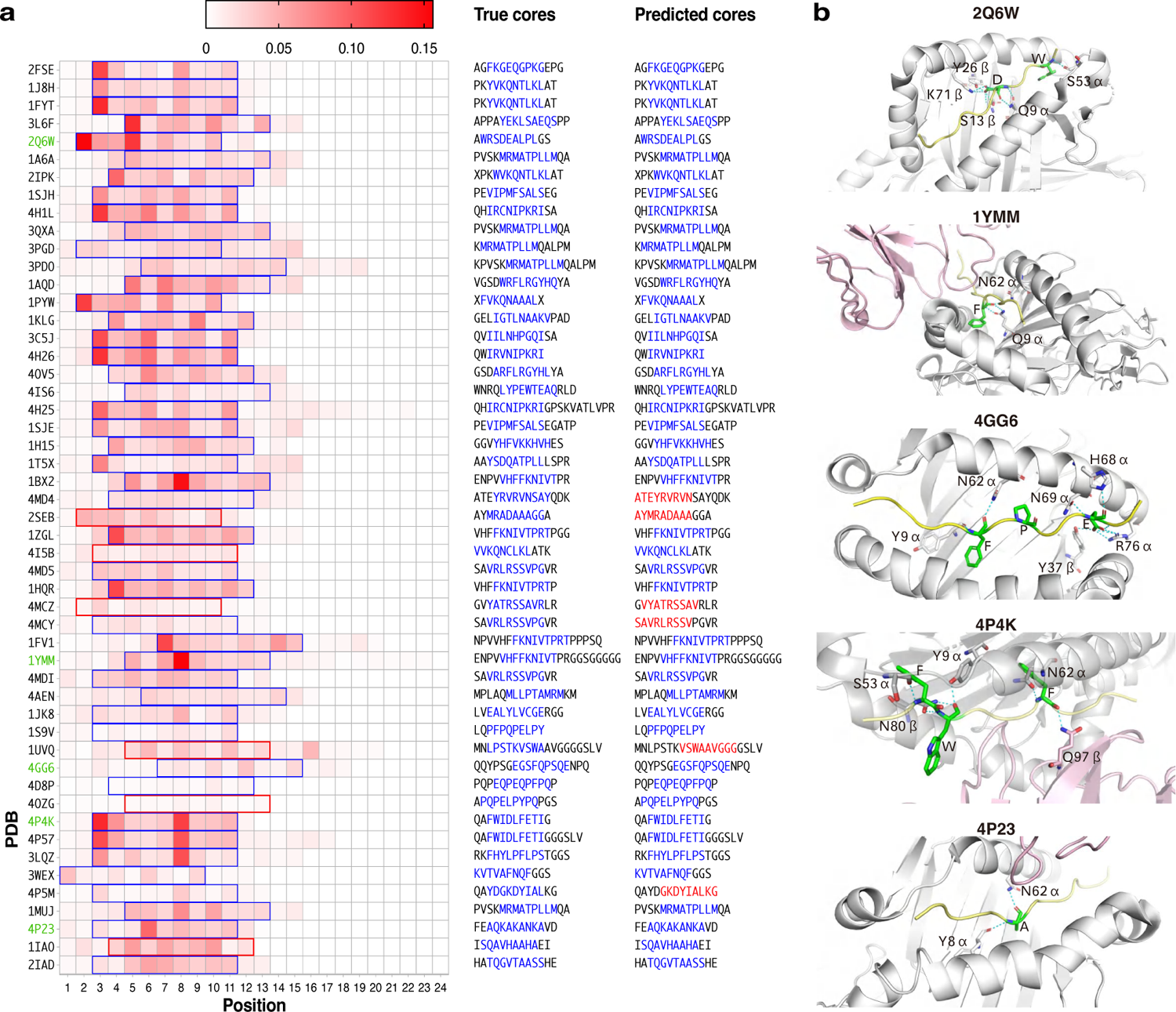
Visualization of ConBoTNet average attention to antigen encodings on BC2015. a. Heatmap visualization of each position of the antigenic peptide sequences of 51 MHC-II peptide complexes on BC2015. Each row in the heatmap represents a visualization of the average attention for each position of the antigen sequence of the complex, and each column denotes the relative position. Each row of the “true cores” column is the corresponding peptide sequence and the binding core is marked in blue. Meanwhile, each row of the “predicted cores” column represents the binding core position predicted by the ConBoTNet method. A correct prediction is colored in blue, whereas an incorrect one is marked red. b. Mark the high attention peptide binding sites (green) on the peptide MHC-II complex structure, along with the polar interactions (blue) with MHC-II (gray) and TCR (pink).

### 3.6. Sequence logo representations

We visualized the binding motifs of MHC-II molecules, identified by ConBoTNet and DeepMHCII, as sequence logos using Seq2Logo v2.0 (Thomsen & Nielsen, 2012). Following the description in You *et al*. (You, et al., 2022), we initially calculated the binding scores for 100,000 random peptides from UniProt (Almagro Armenteros, et al., 2019) in relation to a specific MHC-II molecule. Subsequently, we selected the top 1% of peptides with the highest binding scores to generate the sequence logos under the default settings.

Furthermore, we focused on six MHC-II molecules that demonstrated high predictive accuracy on the BD2023 test set and had detailed motif information in the SYFPEITHI database (Schuler, Nastke, & Stevanović, 2007). The binding groove of the MHC-II molecule is open at both ends, enabling the interaction with peptides of variable lengths. The ends of the binding pocket are called the major anchors (P1, P9) from the N-terminus to the C-terminus, whereas the smaller pockets generate auxiliary anchors (P4, P6, P7) (Ferrante, 2013). **Figure 5** illustrates the sequence logos predicted by two advanced methods, namely ConBoTNet and DeepMHCII, for six MHC-II molecules with diverse anchor types including P1-P4-P6-P9, P1-P4-P7 and P1-P4-P9.

**Figure 5.**
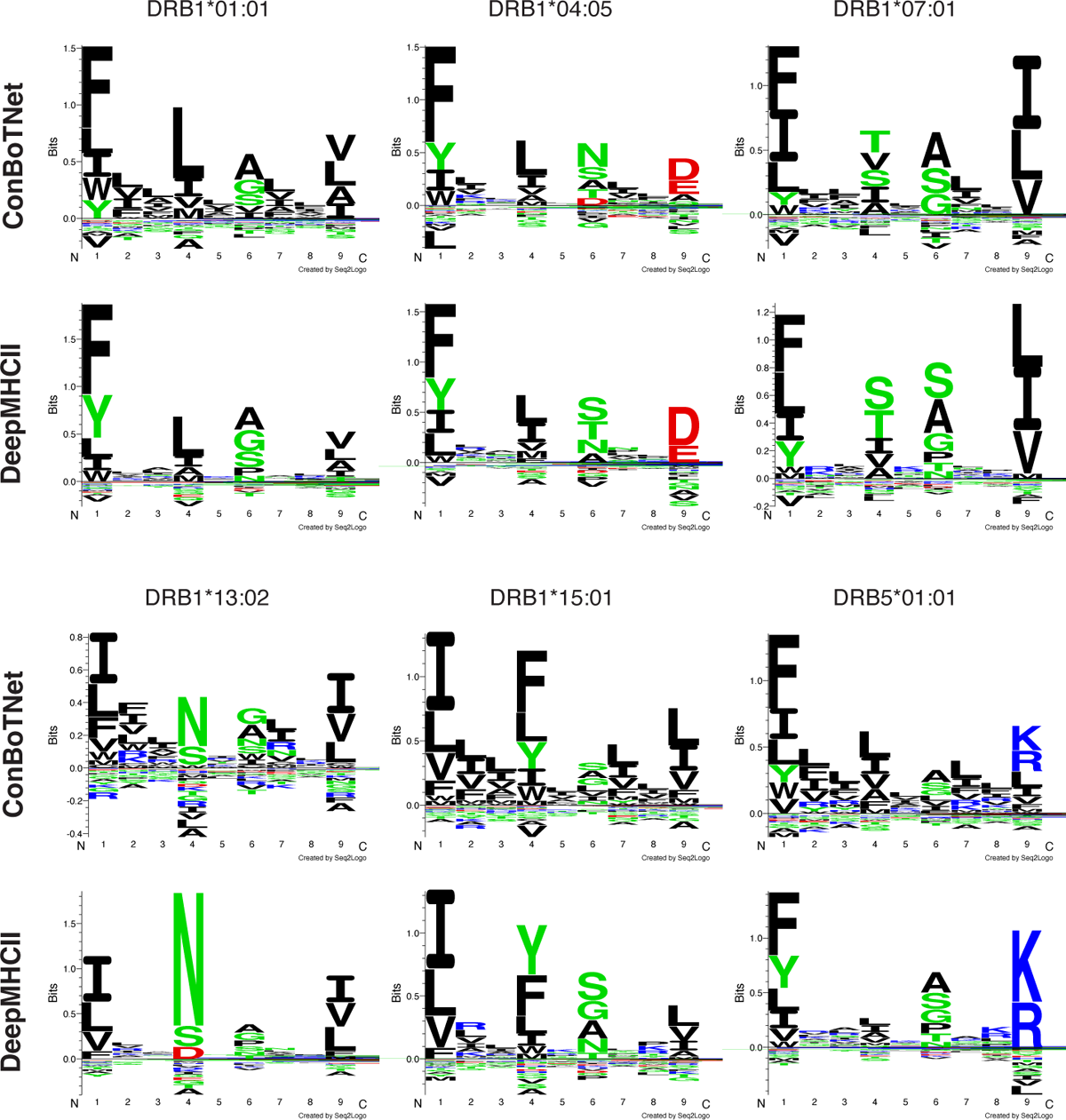
Sequence logo representations by ConBoTNet and DeepMHCII. Each sequence logo has nine positions (pockets) on the horizontal axis. At each position, the total height represents the relative information content of the corresponding position within the motif. Additionally, the height of each letter indicates the frequency of the corresponding amino acid in the position.

Next, we assessed the anchor recognition capabilities of ConBoTNet and DeepMHCII. For the molecules DRB1*01:01, DRB1*04:05, DRB1*07:01, and DRB1*13:02, both methods could identify their anchors, namely P1-P4-P6-P9. However, the distinction between pockets became less pronounced due to the integration of contextual information by the multi-head attention module. Consequently, the anchors identified by ConBoTNet were not as distinct as those provided by DeepMHCII. In the case of DRB1*15:01 with P1-P4-P7 anchors, only ConBoTNet accurately identified P7. Moreover, for DRB5*01:01 with P1-P4-P9 anchors, DeepMHCII failed to identify the correct P4 anchor. Furthermore, we also analyzed the anchor amino acid preferences, whose details are described in **Supplementary Text S5**. Overall, ConBoTNet achieved a more accurate predictive performance for predicting the anchors and amino acid preferences than the competing methods.

### 3.7. Supervised contrastive learning enhances affinity prediction

In this section, we explore the role of supervised contrastive learning (pre-training) in improving affinity prediction models. For more intuitive representations, we output the normalized embeddings through the pre-trained model under 5-fold CV. Next, we use UMAP tool (Becht, et al., 2019), a general-purpose manifold learning and dimensionality reduction algorithm, to visualize these embeddings. **Figure 6** and **Figure S2** show the visualization of the original distribution of the testing data (one-fold) and the embedding visualization of the corresponding testing set under 5-fold CV. Leveraging the labeled data, the supervised contrastive loss encourages the network to pull normalized embeddings from the same class closer, while pushing embeddings from different classes apart. In the pre-training data processing, we hard-divided the data into ten classes according to the affinity value, following **Figure 1e**. Interestingly, the model via supervised contrastive learning (pre-trained stage) can learn a gradient “curve” that varies by affinity value. The robust base model established a solid foundation for the subsequent fine-tuning phase. In addition, the base model can be used to fine-tune MHC-II molecular data with fewer data, enhancing the affinity prediction accuracy for molecules with smaller samples. It also demonstrates why fine-tuning the pre-trained model for only 1-2 epochs can produce excellent prediction accuracy.

**Figure 6.**
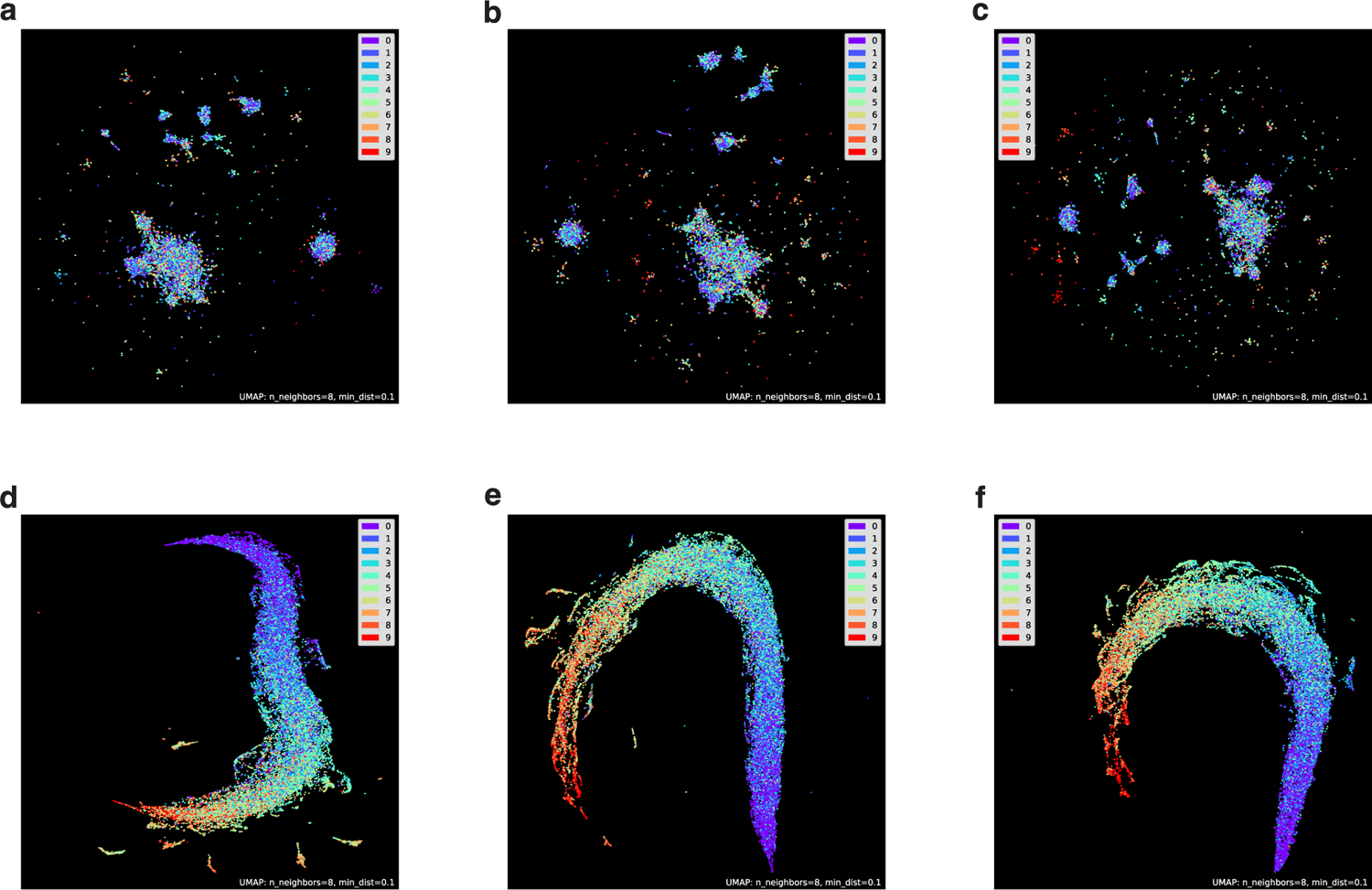
Visualization of the original distribution of the testing data (one-fold) and the embedding visualization of the corresponding testing set under 5-fold CV. a, b, and c respectively correspond to the dimension reduction visualization of the original data of the 1^st^-fold, 2^nd^-fold, and 3^rd^-fold. d, e, and f respectively represent the embedding visualizations learned by the corresponding fold in the pre-training model.

## 4. Conclusion

In this study, we have designed and developed ConBoTNet, a novel deep-learning algorithm for predicting MHC-II peptide binding affinity. The bottleneck transformer module successfully captured the binding core and precise anchor information of peptide MHC-II from multi-source interaction features (including embedding and one-hot encoding), and then learns a robust feature representation based on the similarities and differences between the interaction features through supervised contrastive learning for high-precision affinity prediction.

ConBoTNet demonstrated exceptional performance across multiple benchmark datasets, surpassing competing methods in extensive tests. Furthermore, ConBoTNet identified the binding cores as well as important anchors more accurately than others. In terms of model interpretability, we used the Captum algorithm to explain the influence of binding core and peptides flanking regions on affinity prediction by analyzing the attention weights on peptide sequences; and visually demonstrated how supervised contrastive learning could enhance affinity prediction. All these results highlight the utility and significance of ConBoTNet for relevant biomedical discovery and interpretability. One limitation of our study is that we focused on the peptide-MHC binding prediction rather than T cell epitope identification problem. However, peptide-MHC binding is a prerequisite for T cell immunogenicity, for which multiple studies have shown that MHC peptide binding strength is correlated strongly with peptide immunogenicity (Kim, Bang, Noh, & Choi, 2023). This study provides a useful groundwork for our better understanding of the interaction conformation between T-cell receptors and epitopes. We plan to integrate multivariate prior biological and biomedical knowledge and design highly interpretable model architectures to develop advanced deep learning methods for predicting of T cell receptor-epitope binding specificity (Lu, et al., 2021; Peng, et al., 2023; Racle, et al., 2023) in future work.

## Data and Software Availability

The data and source code for this work are available on GitHub at https://github.com/shenlongchen/conbotnet.

## Supporting Information Available

The Supporting Information is available in the Supplementary file about more details of results. **Text S1**: Performance evaluation metrics; **Text S2**, **Table S3**: Model implementation and hyperparameter settings; **Text S3**, **Figure S1**: Performance on BD2024 dataset; **Text S4**, **Table S1-S2**, **Figure S3**: Ablation experiment; **Text S5**: Analysis of anchor amino acid preferences; **Figure S2**: Visualization.

## Acknowledgments

This work was supported by the National Natural Science Foundation of China (62372234, 62072243), the Postgraduate Research & Practice Innovation Program of Jiangsu Province (KYCX23_0490), the Australia Research Council (LP220200614), both Major and Seed Inter-Disciplinary Research (IDR) projects awarded by Monash University.

## References

Almagro Armenteros, J. J., Tsirigos, K. D., Sønderby, C. K., Petersen, T. N., Winther, O., Brunak, S., von Heijne, G., & Nielsen, H. (2019). SignalP 5.0 improves signal peptide predictions using deep neural networks. Nature biotechnology, 37, 420–423.

Alspach, E., Lussier, D. M., Miceli, A. P., Kizhvatov, I., DuPage, M., Luoma, A. M., Meng, W., Lichti, C. F., Esaulova, E., & Vomund, A. N. (2019). MHC-II neoantigens shape tumour immunity and response to immunotherapy. Nature, 574, 696–701.

Andreatta, M., Trolle, T., Yan, Z., Greenbaum, J. A., Peters, B., & Nielsen, M. (2018). An automated benchmarking platform for MHC class II binding prediction methods. Bioinformatics, 34, 1522–1528.

Becht, E., McInnes, L., Healy, J., Dutertre, C.-A., Kwok, I. W., Ng, L. G., Ginhoux, F., & Newell, E. W. (2019). Dimensionality reduction for visualizing single-cell data using UMAP. Nature biotechnology, 37, 38–44.

Blass, E., & Ott, P. A. (2021). Advances in the development of personalized neoantigen-based therapeutic cancer vaccines. Nature Reviews Clinical Oncology, 18, 215–229.

Chang, S. T., Ghosh, D., Kirschner, D. E., & Linderman, J. J. (2006). Peptide length-based prediction of peptide–MHC class II binding. Bioinformatics, 22, 2761–2767.

Couture, A., Garnier, A., Docagne, F., Boyer, O., Vivien, D., Le-Mauff, B., Latouche, J.-B., & Toutirais, O. (2019). HLA-class II artificial antigen presenting cells in CD4+ T cell-based immunotherapy. Frontiers in immunology, 10, 447508.

Fatima, I., Ahmad, S., Abbasi, S. W., Ashfaq, U. A., Shahid, F., ul Qamar, M. T., Rehman, A., & Allemailem, K. S. (2022). Designing of a multi-epitopes-based peptide vaccine against rift valley fever virus and its validation through integrated computational approaches. Computers in biology and medicine, 141, 105151.

Ferrante, A. (2013). HLA-DM: arbiter conformationis. Immunology, 138, 85–92.

Finotello, F., Rieder, D., Hackl, H., & Trajanoski, Z. (2019). Next-generation computational tools for interrogating cancer immunity. Nature Reviews Genetics, 20, 724–746.

Holland, C. J., Cole, D. K., & Godkin, A. (2013). Re-directing CD4+ T cell responses with the flanking residues of MHC class II-bound peptides: the core is not enough. Frontiers in immunology, 4, 172.

Holling, T. M., Schooten, E., & van Den Elsen, P. J. (2004). Function and regulation of MHC class II molecules in T-lymphocytes: of mice and men. Human immunology, 65, 282–290.

Jensen, K. K., Andreatta, M., Marcatili, P., Buus, S., Greenbaum, J. A., Yan, Z., Sette, A., Peters, B., & Nielsen, M. (2018). Improved methods for predicting peptide binding affinity to MHC class II molecules. Immunology, 154, 394–406.

Jones, E. Y., Fugger, L., Strominger, J. L., & Siebold, C. (2006). MHC class II proteins and disease: a structural perspective. Nature Reviews Immunology, 6, 271–282.

Karosiene, E., Rasmussen, M., Blicher, T., Lund, O., Buus, S., & Nielsen, M. (2013). NetMHCIIpan-3. 0, a common pan-specific MHC class II prediction method including all three human MHC class II isotypes, HLA-DR, HLA-DP and HLA-DQ. Immunogenetics, 65, 711–724.

Kim, J. Y., Bang, H., Noh, S.-J., & Choi, J. K. (2023). DeepNeo: a webserver for predicting immunogenic neoantigens. *Nucleic acids research*, gkad275.

Kisielow, J., Obermair, F.-J., & Kopf, M. (2019). Deciphering CD4+ T cell specificity using novel MHC–TCR chimeric receptors. Nature immunology, 20, 652–662.

Kokhlikyan, N., Miglani, V., Martin, M., Wang, E., Alsallakh, B., Reynolds, J., Melnikov, A., Kliushkina, N., Araya, C., & Yan, S. (2020). Captum: A unified and generic model interpretability library for pytorch. arXiv preprint arXiv:2009.07896.

Liu, B., Shao, Y., & Fu, R. (2021). Current research status of HLA in immune-related diseases. *Immunity*, Inflammation and Disease, 9, 340–350.

Liu, Z., Jin, J., Cui, Y., Xiong, Z., Nasiri, A., Zhao, Y., & Hu, J. (2021). DeepSeqPanII: an interpretable recurrent neural network model with attention mechanism for peptide-HLA class II binding prediction. IEEE/ACM Transactions on Computational Biology and Bioinformatics, 19, 2188–2196.

Lu, T., Zhang, Z., Zhu, J., Wang, Y., Jiang, P., Xiao, X., Bernatchez, C., Heymach, J. V., Gibbons, D. L., & Wang, J. (2021). Deep learning-based prediction of the T cell receptor–antigen binding specificity. Nature machine intelligence, 3, 864–875.

Meraviglia-Crivelli, D., Zheleva, A., Barainka, M., Moreno, B., Villanueva, H., & Pastor, F. (2022). Therapeutic Strategies to Enhance Tumor Antigenicity: Making the Tumor Detectable by the Immune System. Biomedicines, 10, 1842.

Moore, T. V., & Nishimura, M. I. (2020). Improved MHC II epitope prediction—a step towards personalized medicine. Nature Reviews Clinical Oncology, 17, 71–72.

Neefjes, J., Jongsma, M. L., Paul, P., & Bakke, O. (2011). Towards a systems understanding of MHC class I and MHC class II antigen presentation. Nature Reviews Immunology, 11, 823–836.

Peng, X., Lei, Y., Feng, P., Jia, L., Ma, J., Zhao, D., & Zeng, J. (2023). Characterizing the interaction conformation between T-cell receptors and epitopes with deep learning. Nature machine intelligence, 5, 395–407.

Purcell, A. W., Ramarathinam, S. H., & Ternette, N. (2019). Mass spectrometry–based identification of MHC-bound peptides for immunopeptidomics. Nature protocols, 14, 1687–1707.

Racle, J., Guillaume, P., Schmidt, J., Michaux, J., Larabi, A., Lau, K., Perez, M. A., Croce, G., Genolet, R., & Coukos, G. (2023). Machine learning predictions of MHC-II specificities reveal alternative binding mode of class II epitopes. Immunity, 56, 1359–1375. e1313.

Sagan, S. A., Moinfar, Z., Moseley, C. E., Dandekar, R., Spencer, C. M., Verkman, A. S., Ottersen, O. P., Sobel, R. A., Sidney, J., & Sette, A. (2023). T cell deletional tolerance restricts AQP4 but not MOG CNS autoimmunity. Proceedings of the National Academy of Sciences, 120, e2306572120.

Schuler, M. M., Nastke, M.-D., & Stevanović, S. (2007). SYFPEITHI: database for searching and T-cell epitope prediction. Immunoinformatics: Predicting immunogenicity in silico, 75–93.

Soleymani, S., Tavassoli, A., & Housaindokht, M. R. (2022). An overview of progress from empirical to rational design in modern vaccine development, with an emphasis on computational tools and immunoinformatics approaches. Computers in biology and medicine, 140, 105057.

Srinivas, A., Lin, T.-Y., Parmar, N., Shlens, J., Abbeel, P., & Vaswani, A. (2021). Bottleneck transformers for visual recognition. In Proceedings of the IEEE/CVF conference on computer vision and pattern recognition (pp. 16519–16529).

Tadros, D. M., Eggenschwiler, S., Racle, J., & Gfeller, D. (2023). The MHC Motif Atlas: a database of MHC binding specificities and ligands. Nucleic acids research, 51, D428–D437.

Thomsen, M. C. F., & Nielsen, M. (2012). Seq2Logo: a method for construction and visualization of amino acid binding motifs and sequence profiles including sequence weighting, pseudo counts and two-sided representation of amino acid enrichment and depletion. Nucleic acids research, 40, W281–W287.

Traherne, J. (2008). Human MHC architecture and evolution: implications for disease association studies. International journal of immunogenetics, 35, 179–192.

Tsai, S., & Santamaria, P. (2013). MHC class II polymorphisms, autoreactive T-cells, and autoimmunity. Frontiers in immunology, 4, 321.

Venkatesh, G., Grover, A., Srinivasaraghavan, G., & Rao, S. (2020). MHCAttnNet: predicting MHC-peptide bindings for MHC alleles classes I and II using an attention-based deep neural model. Bioinformatics, 36, i399–i406.

Vita, R., Mahajan, S., Overton, J. A., Dhanda, S. K., Martini, S., Cantrell, J. R., Wheeler, D. K., Sette, A., & Peters, B. (2019). The immune epitope database (IEDB): 2018 update. Nucleic acids research, 47, D339–D343.

Wang, Y., Xiang, Y., Xin, V. W., Wang, X.-W., Peng, X.-C., Liu, X.-Q., Wang, D., Li, N., Cheng, J.-T., & Lyv, Y.-N. (2020). Dendritic cell biology and its role in tumor immunotherapy. Journal of hematology & oncology, 13, 1–18.

You, R., Qu, W., Mamitsuka, H., & Zhu, S. (2022). DeepMHCII: a novel binding core-aware deep interaction model for accurate MHC-II peptide binding affinity prediction. Bioinformatics, 38, i220–i228.

Zeng, H., & Gifford, D. K. (2019). Quantification of uncertainty in peptide-MHC binding prediction improves high-affinity peptide selection for therapeutic design. Cell systems, 9, 159–166. e153.

