## Supplementary Material for "Supervised contrastive learning enhances MHC-II peptide binding affinity prediction"

### Supplementary Text

#### Text S1

#### Performance evaluation metrics

In this study, we aimed to evaluate two aspects: the binary classification of the model-predicted MHC-II binding to the peptide and the linear relationship between the predicted affinity values and the true values. Therefore, we primarily employed two commonly used evaluation indicators: the Area under the receiver-operating characteristic curve (AUC) and Pearson Correlation Coefficient (PCC).

AUC is utilized to evaluate the performance of models for binary classification problems. An AUC value approaching 1 signifies a superior model. Also, to classify whether a peptide is bound or not, a threshold of $500nM$ was used (peptides with $IC_{50}$ binding value $< 500nM$ are defined as binders). In other words, converted to the [0,1] interval by the formula mentioned in the **benchmark datasets** section, the predicted value of the binder is $> 0.426$. PCC is commonly employed to evaluate model performance for regression problems, as well as to assess the linear relationship between two variables. PCC values range from -1 to 1, with a value of 1 indicating a perfect positive correlation, -1 representing a perfect negative correlation, and 0 indicating no correlation. As an evaluation index in machine learning, PCC is frequently utilized to quantify the extent of linear correlation between the predicted values of the model and the actual values.

Furthermore, to measure the degree of performance difference between the proposed method and competing methods, we employ the t-test to check the significance of observed performance disparities. Conventionally, a P-value$< 0.05$ is indicative of statistical significance, suggesting a notable performance difference.

**Text S2**

**Model implementation and hyperparameter settings**

The model was implemented using the PyTorch framework (v1.12.1) and trained on a single NVIDIA GeForce RTX 4090 Graphics Card. In the pre-training stage, we adjusted the model parameters to minimize the supervised contrastive loss function via the Adam optimizer. This process was followed by the fine-tuning stage where we optimized the Mean Squared Error (MSE) loss function through the Adam optimizer.

To determine the hyperparameters of the model, we compared the average performance metrics of individual MHC-II molecules under the 5-fold cross-validation (5-fold CV) on the BD2016 dataset. The choice of hyperparameters primarily considered two aspects: sensitivity of the hyperparameters to model performance, and achieving a balance between scale and performance.

Due to constraints in computing resources, we initially filtered out those hyperparameters that significantly impacted performance via the control variable method. Following this, we considered the scale of the model and predictive performance within the limited search space to ultimately determine the final hyperparameter setting. **Table S3** depicts the specific search space of hyperparameters and the finalized hyperparameter settings. This optimal set of hyperparameters was subsequently applied to all experimental comparisons, including leave-one-molecule-out (LOMO) validation, binding core prediction, and independent test.

**Text S3**

**The performance of ConBoTNet and competing methods on the BD2024 dataset**

Furthermore, we downloaded the latest IEDB benchmark dataset from http://tools.iedb.org/auto_bench/mhcii/weekly/ to update our independent test set BD2024 (as of 13-02-2024). NetMHCIIpan-4.1 is currently the best predictor on this platform. We also perform performance comparisons with other methods on this benchmark. The experimental results are shown in **Figure S1**. The ConBoTNet method also demonstrates the best prediction accuracy on both measurement types (binary and IC50).

**Text S4**

**The effectiveness of the bottleneck transformer and supervised contrastive learning**

To test the effectiveness of the bottleneck transformer and supervised contrastive learning in model design, we designed four comparative models (ConBoTNet-Baseline, ConBoTNet-1, ConBoTNet-2 and ConBoTNet) and three sets of ablation experiments. The detailed comparison models are described in **Table S2**. The following three sets of ablation experiments were designed. (1) Investigating the importance of bottleneck transformer modules with location information in feature learning. (2) Study the effect of supervised contrastive learning on model performance. (3) Research the improvement brought about by using the bottleneck transformer modules and supervised contrastive learning at the same time. **Table S2** reports the performance of each compared model on 5-fold CV. Overall, both the bottleneck transformer modules and supervised contrastive learning can improve the affinity prediction accuracy alone, and the combination of the two significantly improves the affinity prediction performance.

**The impact of sequence similarity on the independent dataset**

To explore the robustness of the ConBoTNet method, we used the CD-HIT tool to limit the sequence similarity thresholds (90%, 80%, 70%) between the training set and test set to evaluate the prediction performance of ConBoTNet under different similarity thresholds. As shown in **Figure S3**, the control experiment shows that the prediction performance of ConBoTNet only slightly decreases in accuracy as the sequence similarity threshold reducing.

**Text S5**

**Analysis of anchor amino acid preferences**

Furthermore, with further analysis of anchor amino acid preferences, we observed discrepancies between prediction methods. According to SYFPEITHI, ConBoTNet was able to more clearly recognize the preferred amino acid D in P6 of DRB1*04:05 and S in P9 of DRB1*13:02, while

DeepMHCII failed to do so. In P4 of DRB5*01:01, ConBoTNet successfully identified the preferred amino acid group [LIVAFMQS], whereas DeepMHCII only identified the amino acid group [LIAVMS]. According to SYFPEITHI, P4 allows amino acid Q, which is consistent with the sequence logo predicted by ConBoTNet.

### Supplemental Tables and Figures

**Table S1**. Performance of ConBoTNet with varied pre-training categories under 5-fold CV

| Method | AUC | PCC |
|---|---|---|
| ConBoTNet-5-classes | 0.831 | 0.646 |
| ConBoTNet-8-classes | 0.835 | 0.654 |
| ConBoTNet-10-classes | **0.837** | **0.657** |
| ConBoTNet-14-classes | 0.834 | 0.650 |

**Table S2.** Performance of ConBoTNet and competing methods on single model

| Method | AUC | PCC |
|---|---|---|
| DeepSeqPanII | 0.741 | 0.488 |
| DeepMHCII | 0.821 | 0.628 |
| ConBoTNet-Baseline [a] | 0.822 | 0.627 |
| ConBoTNet-1 [b] | 0.828 | 0.639 |
| ConBoTNet-2 [c] | 0.830 | 0.648 |
| ConBoTNet | **0.837** | **0.657** |

[a] *ConBoTNet-Baseline represents the baseline model, replaces bottleneck transformer modules with residual modules, and no supervised contrastive learning pre-training.*
[b] *ConBoTNet-1 represents the model with bottleneck transformer modules, and no supervised contrastive learning pre-training.*
[c] *ConBoTNet-2 represents the model with supervised contrastive learning pre-training.*

**Table S3.** Hyperparameters of ConBoTNet and the corresponding search space

| Phase | Calibration parameters | Search space | Setting |
|---|---|---|---|
| pre-training | learning rate | {0.001, 0.01, 0.05} | 0.01 |
| | batch size | {256, 512, 1024} | 512 |
| | optimizer | {Adam, SGD} | Adam |
| | dropout probability | {0.25, 0.5} | 0.25 |
| fine-tuning | learning rate | {5e-4, 1e-3, 1e-2} | 1e-3 |
| | batch size | {128, 256, 512} | 256 |
| | optimizer | {Adam, SGD} | Adam |
| | dropout probability | {0.25, 0.5} | 0.25 |

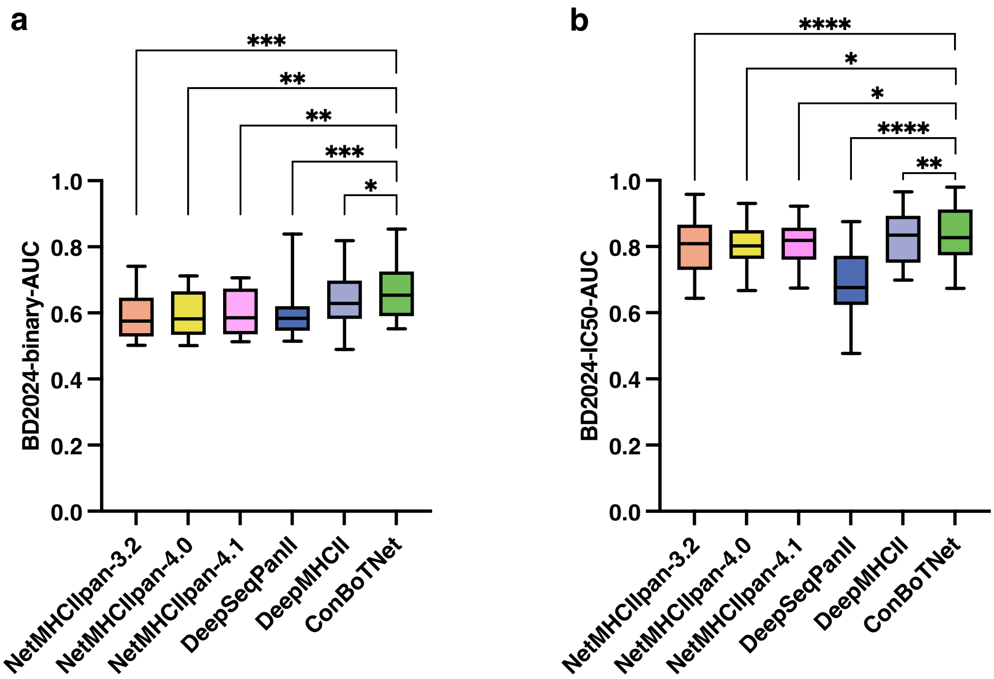


**Figure S1.** The performance of ConBoTNet and other competing methods on the BD2024 dataset. a and b are boxplots of the performance of ConBoTNet and other competing methods on the subset of BD2024 (binary and IC50), respectively.

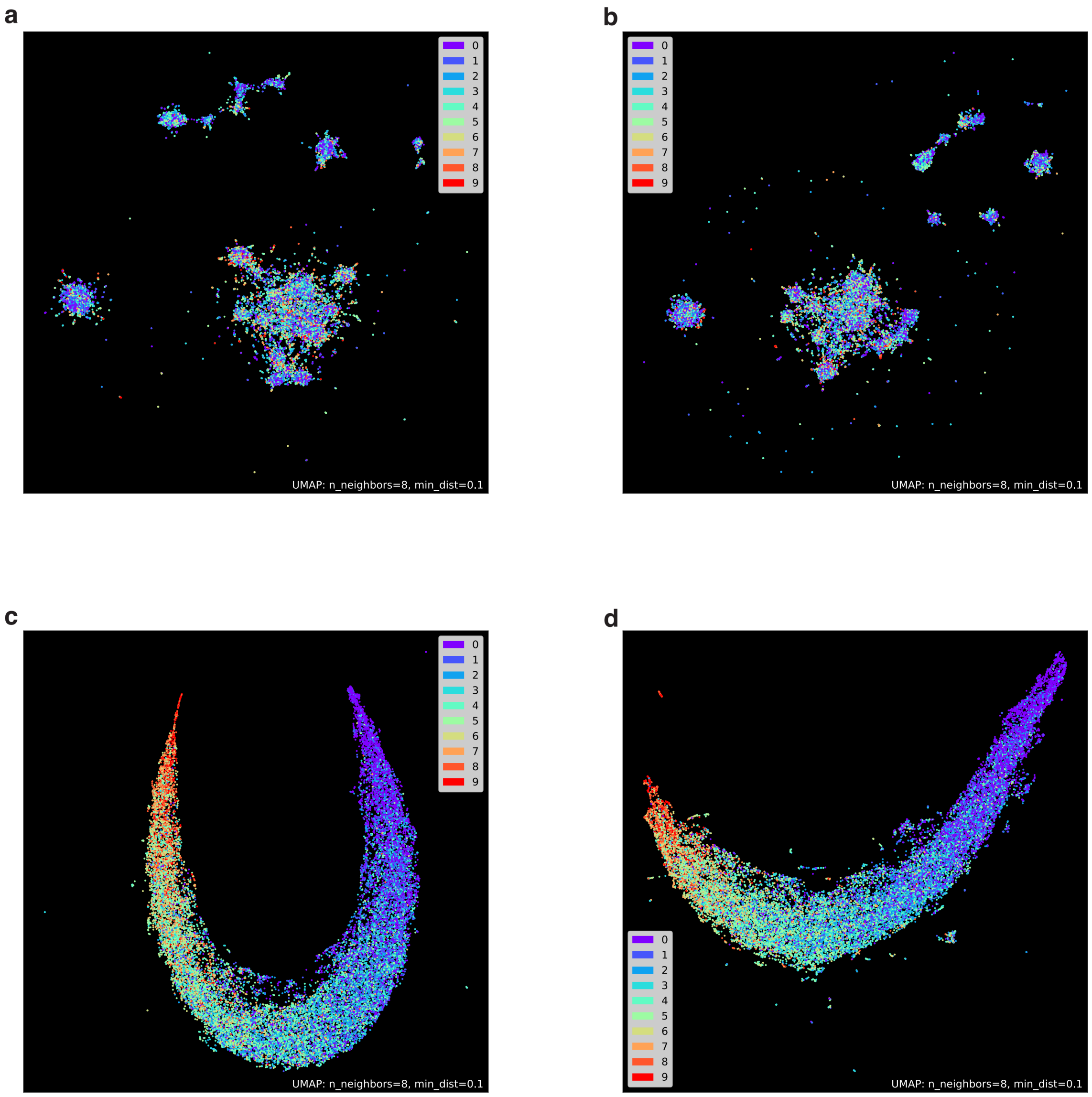


**Figure S2.** The visualization of the original distribution of the testing data (one-fold) and the embedding visualization of the corresponding testing set under 5-fold CV. a and b respectively correspond to the dimension reduction visualization of the original data of the 4th-fold and 5th-fold. c and d respectively represent the embedding visualizations learned by the corresponding fold in the pre-training model.

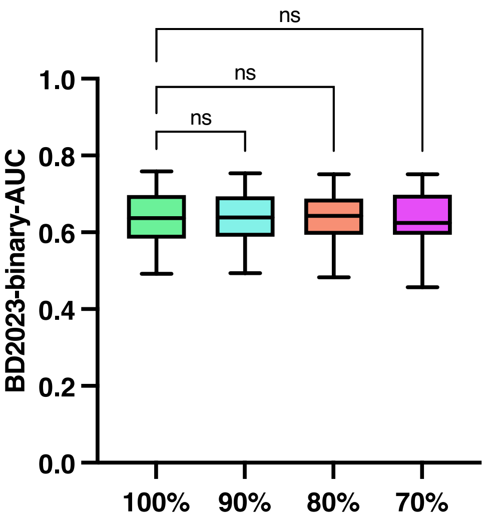


**Figure S3.** The impact of sequence similarity on the BD2024 (binary) subset.
